# Fish Diversity as a Function of Depth and Body Size

**DOI:** 10.1101/208397

**Authors:** Klaus M. Stiefel, Timothy Joseph R. Quimpo

## Abstract

We analyze the number of marine fish species as a function of fish body size and occurrence depth. For this purpose, we analyze the FishBase database. We compare these data to predictions of fish species numbers derived from the neutral theory of biodiversity combined with well-established ecological scaling laws, and measured oceanic biomass data. We consider several variants of these scaling laws, and we find that more large fish species exist compared to the prediction, which is especially true for elasmobranchs, possibly due to their overwhelmingly predatory niches. We find species numbers decreasing with occurrence depth somewhat quicker than our predictions based on the decrease of the number of individuals with depth indicates. This is especially true for the elasmobranchs. This is unsurprising, since the individuals versus depth data did not specifically determine elasmobranch biomass, and since sharks are known to be limited to depths < 3,000 m.

Finally, we discuss how a reduced rate of speciation in larger animals could explain why large species are rare, in spite of the advantages of large body sizes outlined in Cope’s rule.

## Introduction

The world’s oceans are home to an astonishing diversity of fishes, with about 30,000 bony fishes and 900 species of rays and sharks. These fish species range in length from 8 mm (the paedomorphic goby, *Schindleria brevipinguis*, Watson & Walker, 2004), to 12 m (the whale shark *Rhincodon typus*, Fig. 1), and live in ocean zones from the inter-tidal region to the abysmal depths of below 8,000 m (Yancey et al., 2016; maximum depth for sharks ~3,000 m, Priede et al., 2016).

**Figure 1:**
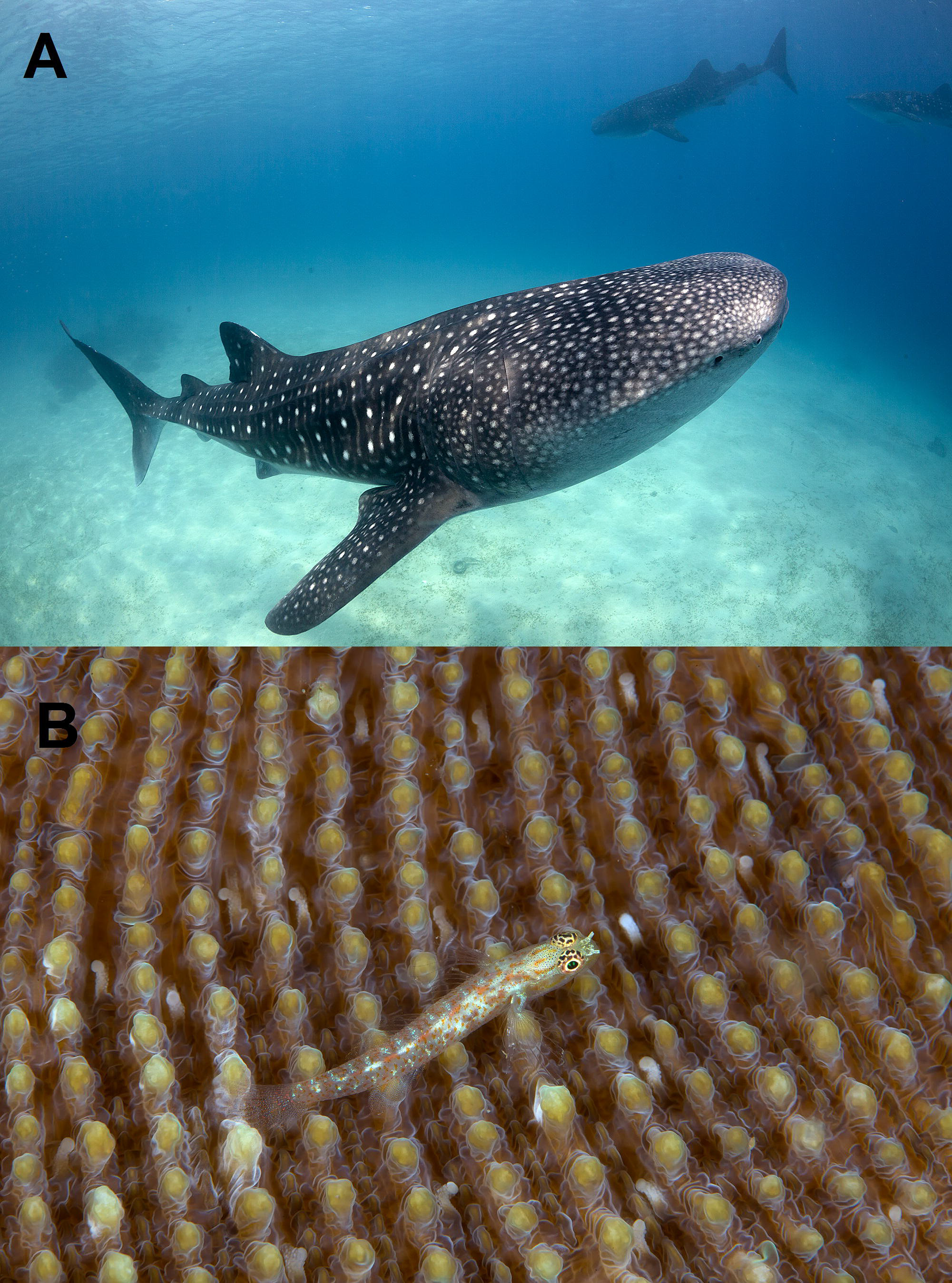
One of the smallest and the largest exant fishes on the planet: A: Whale shark, *Rhincodon typus*, growing up to 12 m and 34 tons. B: Red-spotted pygmy goby, *Eviota albolineata*, attaining a maximum length of only 35 mm. Photographs by K.M.S.

In this study we investigate the distribution of marine fish sizes and occurrence depths with a combination of the analysis of size and depth of 17,163 species with theoretical reasoning. We test if we can predict the size and occurrence depth distributions of marine fishes based on known ecological scaling relationships, measured ocean biomass data and the neutral theory of biodiversity.

Neutral biodiversity theory (Hubbell, 2001; Rosindell et al., 2010; Rosindell et al., 2011; Rosindell et al., 2012) predicts the amount of biodiversity from a number of basic ecological parameters such as population size and speciation rate. Unconcerned with precise biological details unique to each species, it aims to capture grand relationships between fundamental ecological parameters. The theory assumes that two populations of animals with the same number of individuals and the same reproductive rates can be treated similarly in many respects irregardless if the species are insects, mollusks or vertebrates. We apply neutral biodiversity theory to predict fish speciation rates and species numbers as functions of fish size and occurrence depth distributions. Speciation likely occurs in all vertical ocean zones, and in species of all body sizes, but it is not clear how occurrence depth or body size influence the likelihood of speciation. This is the question we address.

## Methods

We analyzed fish size and depth data from FishBase (Froese & Pauly, 2017), using R statistical software (R Core Team, 2017) and the rfishbase package (Boettoger et al., 2012) allowing the importation of FishBase data into the data analysis and plotting language R. For our analysis, we plotted fish species numbers as functions of size and occurrence depth distributions. We then tested if we can successfully predict the observed frequencies based on the neutral biodiversity theory and ecological scaling relationships explained below.

FishBase has been used as the basis for a number of insightful analyses, like of the colonization of the deep sea by different fish families (Priede and Froese, 2013) and of the socio-economic & ecological patterns of biodiversity loss world-wide (Clausen & York, 2007). Naturally, a data base of all fishes will not be complete, since there are still new fish species described every year, and not all parameters will be known with good confidence for every species. However, the data contained in it is vast, and we believe it is an excellent tool for the analysis of evolutionary trends involving many species.

To test for trends in the missing data which could influence our analysis, we plotted the number and percent of species with missing size (Fig. 2) and depth data (Fig. 3) versus the average depth for all 39 analyzed actinoperygian orders and 13 analyzed chondrychtian orders. The percentage of coverage for different orders was between 0% and 100% for minimum/maximum depths, and between 22% and 100% for fish lengths. No trends were apparent, with an even distribution of the coverage values between the extrema. The by far most species-rich order, the Perciformes, had a coverage of 32% and 26% for the minimum/maximum depths, and of 79% for the lengths. In summary, our data-set contained the parameters we analyzed for a significant fraction of all known fish species, with no apparent trends in the omissions.

**Figure 2:**
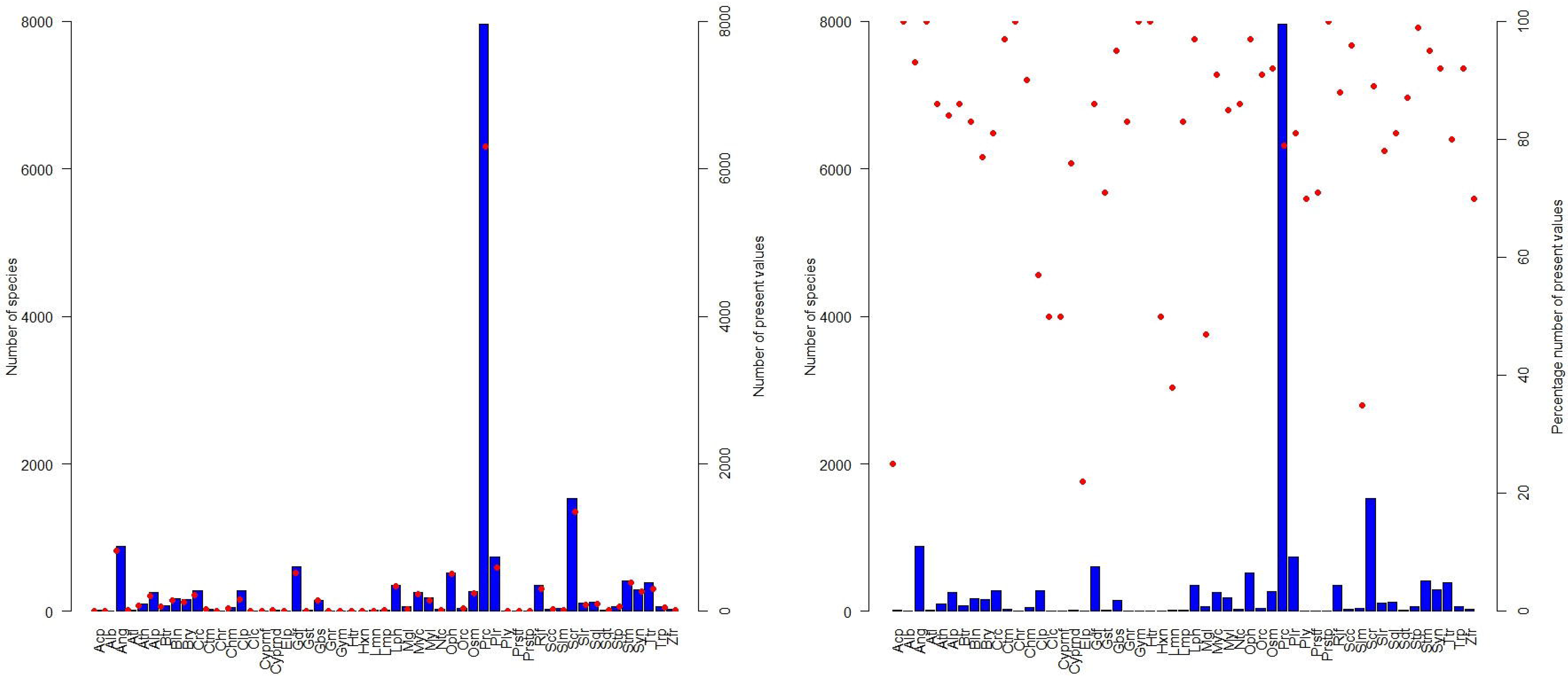
Fish orders, and the number of the species they contain (blue bars) and entries in FishBase (red dots) for length, given in A) number of species with an entry for length and B) the percentage of species with an entry.

**Figure 3:**
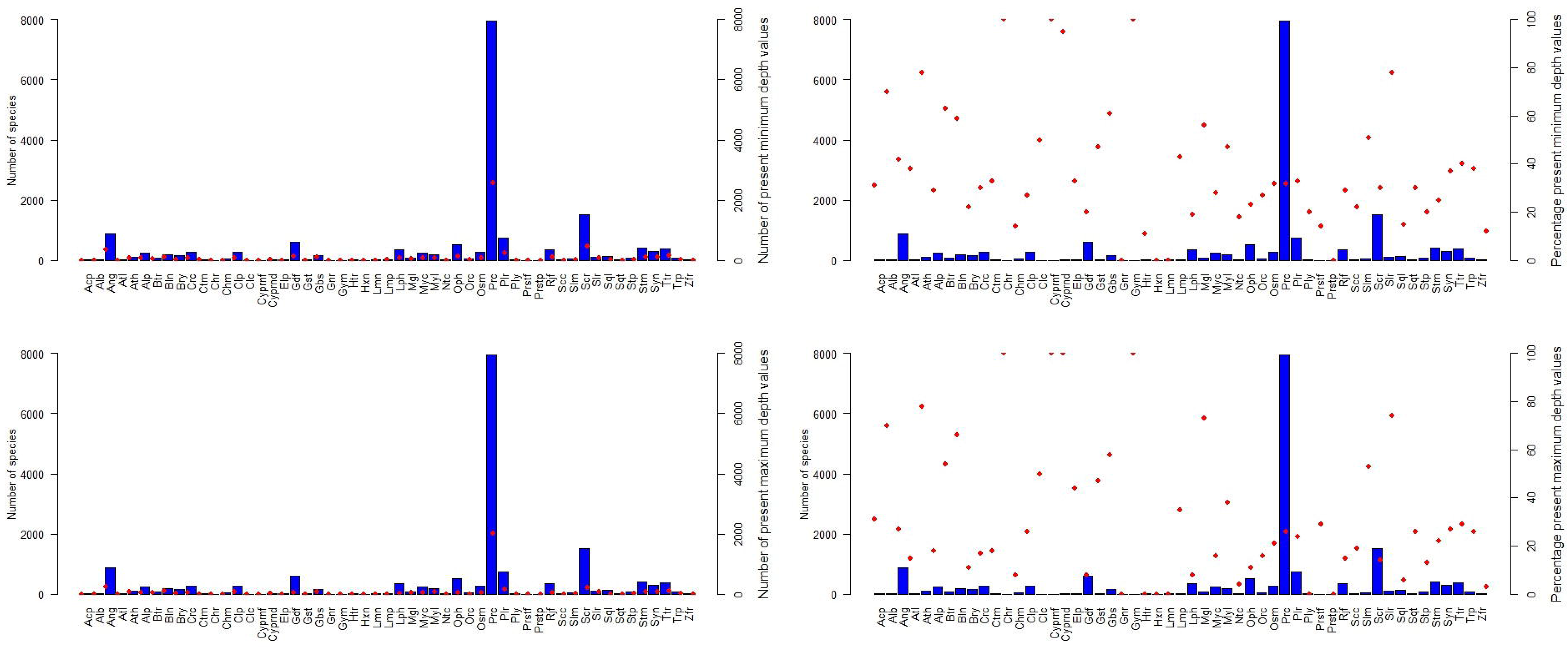
Fish orders, and the number of the species they contain (blue bars) and entries in FishBase (red dots) for A) minimum depth, B) maximum depth, C) percentage of species with an entry for minimum depth and D) percentage with an entry for maximum depth.

In this study we are pooling species from the intertidal zone with those from the deep sea, filter-feeders with herbivores and predators, and broadcast spawners with nest-building fishes. Naturally, speciation in these animals is not comparable in many ways. However, the neutral theory of biodiversity allows a comparison over many very different species. We take advantage of this theoretical framework to interpret large-scale patterns of fish diversity in the ocean.

## Results

We will now first derive predictions for fish species number based on fish size and occurrence depth. We will do this by combining the neutral biodiversity theory and well-established scaling laws. We then test these predictions against FishBase data (for a schematic explanation of our reasoning see Fig. 4).

**Figure 4:**
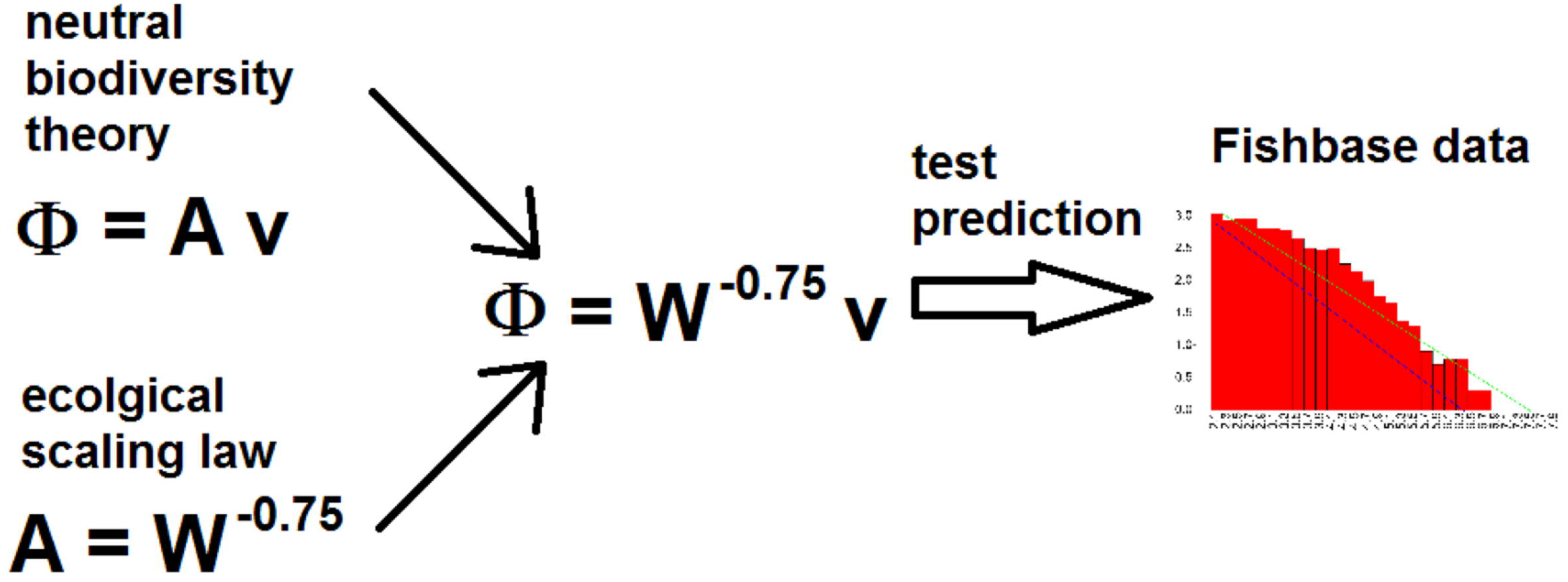
Schematic explanation of the reasoning used in this study. An expression of species number as a function of population size from neutral biodiversity theory together with ecological scaling laws provides predictions for fish species numbers as functions of fish size and occurrence depths. The predictions derived in this way are tested against data from FishBase.

**Figure 5:**
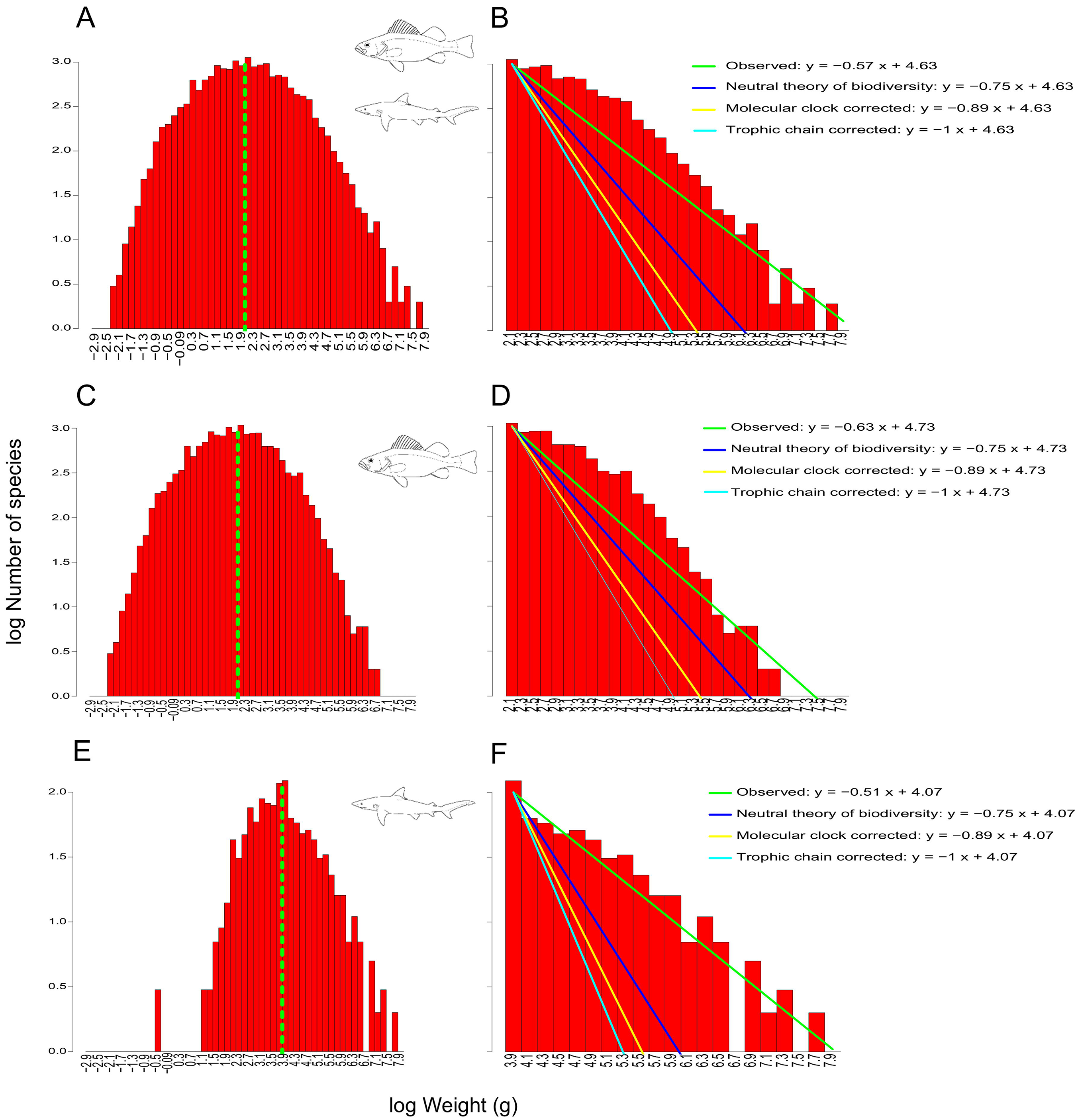
Histogram of fish species weights, showing A) all fishes, B) bony fishes and C) cartilaginous fishes. The best fit to the data from the peak (the size class with most species) on is shown in green, the predicted slope (with an intercept identical to the fit) in blue. A fit corrected for the slower molecular clock at larger sizes is shown in yellow, corrected for a loss of energy towards higher tropic levels in turquoise.

**Figure 6:**
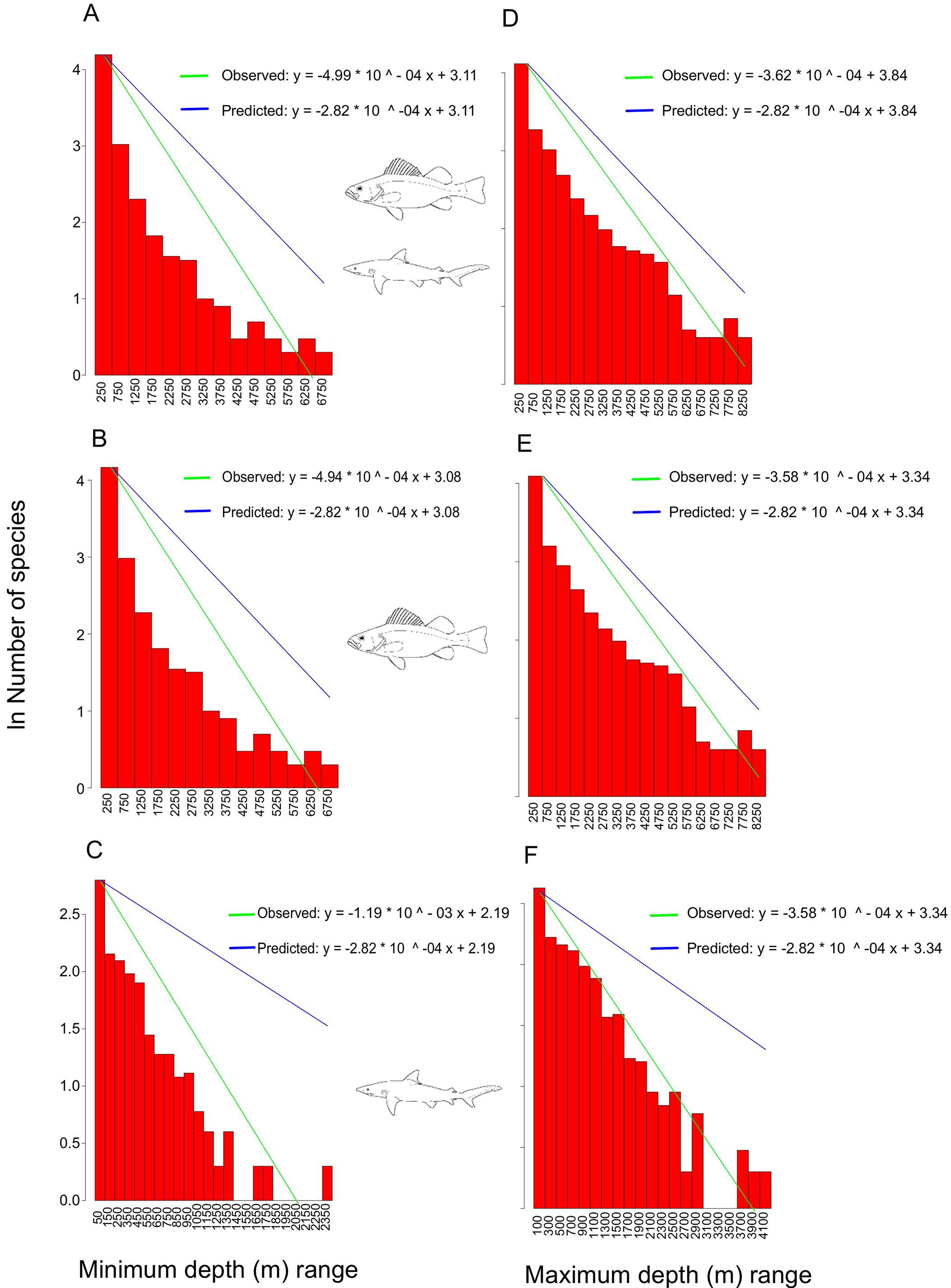
Histogram for minimum occurrence depths for A) all fishes, C) bony fishes and E) cartilaginous fishes, and maximum occurrence depths of B) all fishes, D) bony fishes and F) cartilaginous fishes. The best fit to the data is shown in green, the prediction derived from the decreasing recorded number of individuals (Wei et al., 2010) is shown in blue.

A fundamental prediction of neutral biodiversity theory is that the likelihood of speciation follows a simple relationship:

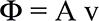

Which is that the speciation (Φ, new species per generation) is a product of the abundance of a meta-population (A, individuals) multiplied with a speciation rate (v, per individual per generation, alternatively per time, or per unit of metabolism). The debate if mutation/speciation rate is best described as per generation, or per time, or metabolism is not settled; likely a mutation/speciation rate per time, is correct (Martin & Palumbi, 1993; Palumbi, 1994). The mutation/speciation rate seems to be roughly a function of total accumulated life-time metabolism (Speakman, 2005), and can be assumed to be a constant or negatively proportional to body-size (see below).

With the relationship stated above the neutral theory of biodiversity provides us with an expression for the *amount of speciation occurring as a function of the population size of a species*. If we assume a constant extinction rate (in the absence of mass extinctions, Raup, 1991; Sepkoski, 1992, 1996) we can relate the population size to the number of species of animals with that particular population size.

As a next step, we need to find *estimates of population sizes as functions of fish body sizes and occurrence depths*. With these estimates we can then *predict the number of fish species as a function of fish body sizes and occurrence depths*. Finally, we can *test these predictions with data covering a large fraction of all known fish species*, provided by FishBase (Fig. 4).

### Species Number/Body Size Relationship

There is a large body of theoretical work on the scaling of body size (W, kg), generation time (G, years), metabolism rate (E, Joule time^−1^) and abundance (A, number of individuals) across different species. Roughly, with some caveats, the following holds (Speakman, 2005):

G = W^0.3^ (larger animals live longer)
E = W^0.7^ (larger animals use more energy)
A = W^−0.75^ (larger animals are less abundant, Marquet et al., 2005)

These scaling laws are not precise expressions, rather they are correct by an order of magnitude. However they explain variations in size, life duration and abundance over many more orders of magnitude. These scaling laws hold better within than between phylogenetic groups, for example mammals and birds have somewhat different exponents describing energy use as a function of size (Speakman, 2005). The scaling laws also hold better in some groups than in others. While these scaling laws have to be interpreted with caution, at least the relationships between body size, longevity and metabolism likely have a biochemical basis (Speakman, 2005), and hence a mechanistic foundation.

For a prediction of the number of fish species as a function of their body size we combine the aforementioned relationships with the expression for speciation:

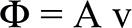

*and*

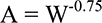

*thus*

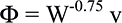

Hence, the number of species of fish of a certain size is related to the size (weight) raised to the negative power of 0.75. This is the scaling relationship believed to generally hold, based on energy availability (Marquet et al., 2005). A number of studies have, however, suggested modifications of this predictions:

In size-structured food webs the energy flow between primary producers, small prey and large predator can be limited, resulting in a steeper slope of the size/abundance relationship (Rossberg et al., 2008; Blanchard et al., 2008). Slopes around −1, as steep as −1.2 have been reported (Jennings & Mackinson, 2003).

Another modified prediction takes into account that the speciation rate v is slightly decreased by a decreased mutation rate in larger animals. The nucleotide replacement rate is 9 times lower per 10^5^ increase in body size, corresponding to a 9 10^−8^-fold mutation rate decrease per additional g body mass (Martin & Palumbi 1992). Hence we can correct the mutation rate v:

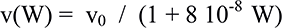

and we get a corrected expression for the speciation rate

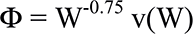

which is

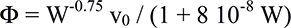

and corresponds to an increase of the slope by a factor of 1/log(8 10^−8^) = 0.14 in a plot with a logarithmic x-axis (like the species number/size histograms). The prediction, after a correction for a decreased mutation rate in larger animals, would be about:

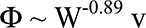

We next test these predictions against empirical data from FishBase. We can estimate the average body weight of a fish species, W, from fish length by using the relationship:

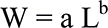

The parameters *a* and *b* depend on the body shape of a fish and are available in the FishBase database (Froese, 2006; Froese et al., 2013). When we plot a histogram of the number of species per fish weight class we see that the distribution peaks at a small size (histogram bin center 10^2.1^ = 126 g). A significant fraction (9,140 or 53%) of fish species attains maximum weights below that. There are several possible reasons which limit the minimum size of a vertebrate to about 8 mm. These include the small number of eggs such a small organism can produce (Depczynski & Bellwood,2005) or the minimum number of individual cells of each cell class necessary for a functioning animal.

Interestingly, several lineages of fishes (Watson & Walker, 2004; Rüber et al., 2007; Kottelat et al.,2006), amphibians (Das & Haas, 2010) and reptiles (Hedges & Thomas, 2001) have all converged to a minimum size just below one centimeter. These constrains likely limit the evolution of smaller vertebrate species even somewhat above the absolute size minimum.

Hence we made the decision to only test our data against the declining (rightward) slope of the size/species number distribution. We fit a linear regression to the linear/log (species number/size) histogram. We found slopes of −0.57 for all fishes, −0.63 for bony fishes and −0.51 for sharks and rays. Thus, the data most agree with a prediction based on the neutral theory of biodiversity (slope −0.75), lying somewhat below this prediction. The lower slope for elasmobranchs possibly indicates that in this class of fishes, with their mainly predatory niches, a stronger selective pressure for larger body sizes exists.

The data agrees worse with a prediction corrected for a lower mutation (speciation) rate of larger animals (Slope ~ −0.89). The species numbers as a function of body size also do not agree with the prediction of steeper slopes found in size-structured marine ecosystems with a limited energy flow to larger animals (Slope < −1). This could have two causes: 1. that the majority of ocean ecosystems is different from the structured communities studied, with an unabated energy flow to animals at larger sizes. Larger animals might be more abundant. In fact, some studies indicate an over-proportional energy use of large animals. 2. Alternatively, speciation might deviate from the proportionality described in the neutral theory of biodiversity (more speciation at large animal sizes than predicted). We can not distinguish these cases at present.

### Species Number/Depth Relationship

We also predicted the number of species as function of occurrence depth. A recent study has explored the depth/species number relationship in great detail (Priede and Froese, 2013) and noticed a slope similar to the slope of oceanic biomass as a function of depth. Data about the marine biomass and number of individual animals as a function of ocean depth is available (Wei et al., 2010), and gives an estimate of the fish population size A (individuals) as a function of depth (z):

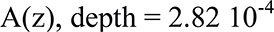

Which is the slope of the decrease of the number of individuals with depth of the marine “megafauna” given in Wei et al., 2010, which includes all fishes and non-microscopic invertebrates.

From this depth/species number estimate we can calculate the speciation rate as a function of depth:

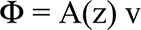

with constant v. Again, with constant extinction rates Φ should translate into species numbers.

We plotted the number of species as a function of and minimal and maximum occurrence depth, analyzing 17,163 species from the FishBase database. The slopes of the fits to the histograms for all fishes, bony fishes and sharks & rays were 4.99 10^−4^, 4.94 10^−4^ and 1.19 10^−3^, respectively for the minimum depth, and 3.62 10^−4^, 3.58 10^−4^ and 6.37 10^−4^ for the maximum depth.

These slopes are somewhat higher than the predictions derived from the neutral theory of biodiversity and the measured (Wei et al., 2010) number of individuals as a function of depth. As expected from the lack of sharks at deeper depths (Priede et al., 2006), the slopes were especially steep for elasmobranch fishes.

As in the case of the lower mutation (and hence speciation) rate of larger animals for which we corrected in the first part of the study, the mutation rate will likely be lower at deeper depths, due to colder ambient temperatures. Since mutation rate is roughly proportional to the metabolic rate, and the metabolic rate roughly doubles with a 10°C increase in temperature (q_10_ ~ 2), the mutation rate will be lower at colder temperatures. Unfortunately, there is no ubiquitous expression for the decrease in water temperature with ocean depth, since different oceans differ in topology, currents, upwellings and other physical factors. While we can’t quantitatively adjust our predictions, we can be certain that deeper parts of the ocean will be cooler (with rare exceptions) and the mutation rates will be lower. This depth-temperature relationship might play a role in explaining the lower than expected species number at depth.

## Discussion

We analyzed the number of marine fish species as a function of their size and occurrence depth. We conducted these analysis for all fishes and bony fishes and sharks separately. When relating species number to size, we found that it decreases with slopes smaller than predicted by neutral biodiversity theory and ecological scaling rules (more large fishes than predicted). Modifications of the prediction based on a lower mutation rate of large animals or on an energy-flow limiting food web made the predictions worse, not better. Elasmobranch species were especially found to be larger than predicted.

When relating species number to occurrence depth, we found that the slopes were higher than predicted by the neutral biodiversity theory and the measured drop in the number of individual marine animals with depth (fewer species at depth). This is possibly due to colder temperatures, and hence lower mutation rates at depth. Unfortunately this drop in temperature is hard to quantify.

In general, the data covering a significant part of all known marine fishes agrees with the predictions by the neutral biodiversity theory, and scaling laws/animal counts at depth moderately well. As has been pointed out previously (Rosindell et al., 2010), the neutral theory of biodiversity can at lest serve as a good starting point for any ecological investigation. If a deviation from this initial prediction is found, the search for reasons for the predictions will give insights into the investigated species and ecosystems. Here, we speculate that the colder temperatures, and hence the reduced metabolism and mutation rates, cause a reduced number of fish species at depth. We also speculate that the predatory life-style of sharks causes a larger deviation of their species number from the initial prediction; and that the general under-estimation of the number of large species is due to the many advantages conferred by large body size - as stated in Cope’s rule.

Cope’s rule postulates that due to the several advantage of being large many animal lineages produce large or very large species later in their evolutionary history (Hone & Benton, 2005; Hone et al., 2005). These advantages include a reduced predation pressure against large animals, improved digestion with long intestines, and more efficient thermoregulation. Indeed, many lineages like dinosaurs (Hone et al., 2005), and cetaceans have produced very large species, while the rule did not take effect in other lineages such as bivalves (Jablonski, 1997). One question is why, with the advantages of large size, not the majority of species are large. The reasoning followed here connects speciation rates to body size. Speciation is slow at large sizes, with a lower abundance of large animals, and large species are removed by extinction at the same rate as more numerous small species (Fig. 7). This suggests a process where small species speciate fast and give rise to many new species. Some of these species grow in size, according to Cope’s rule. However, these big species then show slow speciation, which in the presence of constant extinctions can lead to an extinction of the complete lineage. In this way, despite the advantages of large body size, the steady-state species number versus species size distribution is skewed towards small species. A similar scenario, involving extinction events reducing the number of large species is suggested Arnold et al. (1995, in their investigation of planktonic foraminifera).

**Figure 7:**
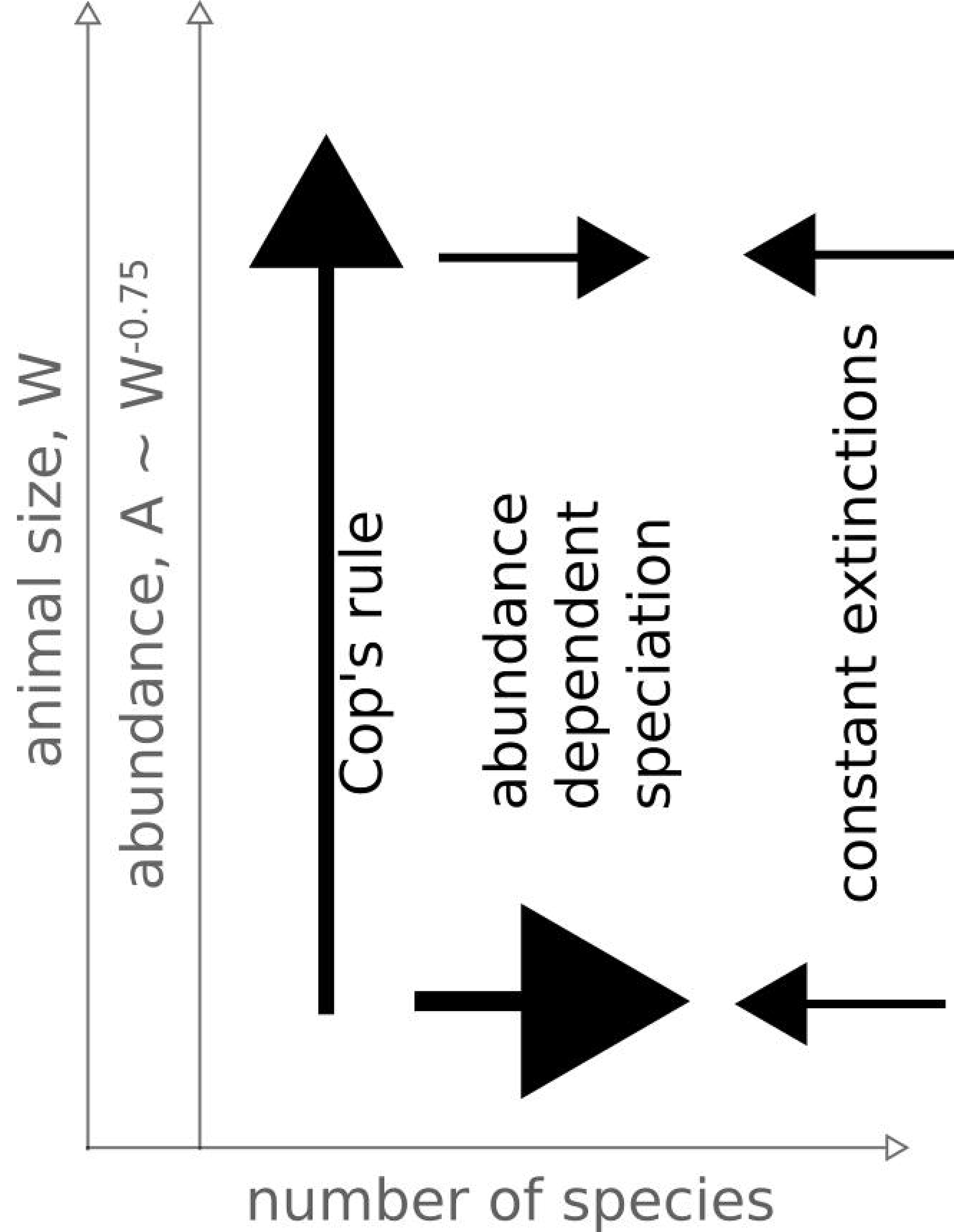
Schematic explanation of the relationship between size-dependent and hence abundance dependent speciation and Cop’s rule.

